# Co-Designing Research Priorities in Developmental Neuroscience: A Place-Based Participatory Priority-Setting Study

**DOI:** 10.64898/2026.01.06.697951

**Authors:** Katherine Hiley, David Ryan, Layla Kouara, Zarina Mirza, Mark Mon-Williams, Faisal Mushtaq

## Abstract

Children and adolescents, the primary participants in developmental neuroscience research, are rarely consulted or included in the research design process. By neglecting their perspectives and needs, we risk limiting the relevance and effectiveness of translational research that may emerge from this work. Here, we report on a process of co-design with adolescents to inform the generation of a new population neuroimaging research programme. We employed a two-study community-based participatory approach and used mixed methods to identify the research priorities of this community. In the first study, 79 secondary school students from four schools were introduced to fundamental neuroscience concepts during five one-hour workshops. In groups, they then designed research studies, centred around topics they deemed important, and were asked to present their ideas through poster presentations, role playing as neuroscientists. Reflexive thematic analysis was conducted on these posters, extracting ten key themes. In the second study, 376 students from four different schools, ranked these ten themes in order of importance. Mental health and stress consistently emerged as the highest priorities. Further exploratory analysis revealed that the priority for exploring everyday routines and understanding anti-social behaviours varied across schools and age groups, while the importance of mental health varied significantly between genders. This work provides an illustration of how to implement place-based participatory approaches at the early stages of a developmental neuroscience research project to maximise inclusivity and potential downstream translation and impact.

## Introduction

Co-design in health research is a collaborative approach that involves researchers, healthcare professionals, and community members as equal partners throughout a project (NIHR, 2024). From initial planning through to generating new insights, all partners share authority and decision-making. This process, now regularly employed in community-based studies, ensures that a range of perspectives are included, increasing the relevance and practical value of findings. By incorporating stakeholder insights at every stage, co-design can help align scientific goals with community needs and increase the chances of producing outcomes that have real-world impact (Tembo et al., 2021). Such approaches have also shown value in global health contexts (National Institute for Health and Care Research Global Health Research Unit on Global Surgery, 2023).

While its advantages are clear, co-design is rarely employed in developmental neuroscience. This lack of community engagement can be seen as part of a broader concern about the limited diversity in neuroimaging research, where WEIRD (Western, European, Industrialised, Rich, Democratic) populations are disproportionately represented (Ricard et al., 2023; Webb et al., 2022). Recent work has begun to explore how community-based participatory research principles may be applied within developmental neuroscience settings (La Scala et al., 2023), although systematic documentation of participatory approaches within the field remains limited. This follows a decade of calls for stronger cultural competency, improved transparency, and the active involvement of marginalised groups in all phases of research (Mikesell et al., 2013), highlighting that more needs to be done to ensure research agendas truly represent and include the voices of groups that are often overlooked. By integrating CBPR principles into the process, from the earliest stages of planning and design, researchers can build genuine partnerships with communities. This approach can not only strengthen collaboration but also help researchers develop cultural awareness and understanding (Toenders et al., 2024). In the long run, it could also deliver a step toward addressing the longstanding inequalities that continue to affect developmental neuroscience research.

Examples of such involvement frequently uncover possibilities that might be overlooked when adults work alone (Abraczinskas & Zarrett, 2020), and demonstrate how co-design can shape the research process (Clark et al., 2022). One such example comes from the Kids Interaction and NeuroDevelopment (KIND) lab, where informal conversations with pre-adolescent Latina girls revealed an association between minority ethnic status and internalised social anxiety. As a result, the lab shifted its focus to explore how Latina mothers’ discussions about ethnic and racial identity influence their children’s emotional expression and regulation and incorporated participant feedback to alter themes and questions within their surveys and recruitment strategies to better capture mental health outcomes in Latina youths (La Scala et al., 2023)..

Adolescents, while clearly central for developmental neuroscience research, have been largely excluded from influencing research agendas. Adolescence is also a time marked by extensive brain development in regions including the prefrontal cortex and limbic system, areas associated with mental health vulnerabilities (Paus et al., 2008). The rising prevalence of mental health challenges among adolescents, exacerbated by academic pressures, social media further emphasises the potential value in engaging this group (Orben & Przybylski, 2019; Pierce et al., 2020; Twenge et al., 2018).

Involving adolescents in co-design may also address a key developmental need to contribute, which is often overlooked (Fuligni, 2019), while also ensuring that study goals and methods are directly relevant to their lived experiences (Ryan et al., 2024). Adolescents who participate in research are reported to gain skills and a sense of purpose as they see their input shape real decisions, echoing findings that co-produced youth involvement can build confidence, insight, and shared ownership (Pavarini et al., 2019). Evidence from evaluations of young people’s advisory groups shows similar benefits, with adolescents reporting belonging, skill development, and confidence, while researchers gain perspective and refine their methods through reciprocal learning (Brady et al., 2023). Such authentic input can strengthen research design and foster trust and enthusiasm between young people and researchers (Ryan et al., 2024; Warraitch et al., 2024).

In youth settings, adolescent perspectives become especially important for designing research with greater resonance, as local knowledge and lived experiences enhance the depth and real-world applicability of findings (Marschall, 2004). Collaborating with adolescents in this way can align with evidence that combining institutional resources with active stakeholder participation strengthens the validity and policy relevance of research (Ostrom, 1996). The Born in Bradford “Age of Wonder” programme (Shire et al., 2024), a seven-year project capturing the journey through adolescence and adulthood for all teenagers in the UK city of Bradford using health, social and education data, is a living example of these benefits. The BiB programme has shown how young people’s involvement can not only improve research quality but can also facilitate skill development and empowerment (Ryan et al., 2024). The project employed iterative “reflection circles” and qualitative feedback from “co-researcher” participants to refine critical thinking and problem-solving skills, as they were actively involved in addressing real-world issues and making decisions that directly impacted research outcomes. In these sessions, participants described how much they felt responsible for steering research decisions (ownership) and their confidence in shaping outcomes (self-efficacy). These perceptions were then checked across multiple inquiry cycles to capture changes over time, if, for instance, participants reported growing comfort in contributing ideas and observed their suggestions being integrated into the research design. This cyclical approach provided direct, evidence-based insights into how and why co-design enhanced adolescents’ sense of agency.

Here, building on this work, we report on a structured, multi-study community-based participatory approach to co-develop research priorities for a new population neuroimaging programme with adolescents. In this paper, we describe our work as co-design rather than co-production. This distinction is intentional. Co-production typically implies that stakeholders are involved across all stages of the research cycle, from identifying priorities, shaping methods, collecting and analysing data, through to dissemination and implementation (NIHR, 2024). Here, we use ‘co-design’ to refer to participatory processes where community members hold meaningful influence over research priorities.

Our approach involved interactive workshops that introduced adolescents to foundational neuroscience concepts and engaged them in shaping research priorities through creative, collaborative activities. We had two aims, to: (1) illustrate a methodological framework for participatory priority-setting in developmental neuroscience; and (2) report the specific research priorities that adolescents themselves identified as important.

## General Methodology

The neighbouring cities of Bradford and Leeds in West Yorkshire reflect the UK’s diverse social and economic landscape. Bradford is England’s fifth most deprived district, with 40% of children living in poverty higher than the regional and national averages (City of Bradford Metropolitan District Council, 2023). Socioeconomic disadvantage is associated with reduced access to educational and health resources and increased risk of poorer cognitive and mental health outcomes in young people (Farah, 2017; Reiss, 2013). Some Bradford schools, however, have received national recognition for innovative, community-based education programmes focused on inclusion and resilience (Ofsted, 2021). Leeds has a more varied economy, but still faces pockets of severe deprivation, especially in former industrial areas (Leeds City Council, 2023). Many schools in these neighbourhoods contend with similar challenges, including food insecurity, housing instability, and limited access to early intervention services, while more affluent districts benefit from ample resources and some of the highest academic attainment in the country (Office for Students, 2022). These contrasts highlight the importance of place-based approaches to education and child development (Cardenas-Iniguez et al., 2024; Thijssen & Van Dooren, 2016). Building on local infrastructure and existing partnerships, we engaged adolescents to explore their priorities for a new population neuroscience research programme.

A two-study mixed-methods design was implemented. Participants for both studies were recruited through school networks established by the aforementioned Born in Bradford project (Shire et al., 2024), the Centre for Applied Education Research (CAER) in Bradford and the Child Health Outcomes Research at Leeds (CHORAL) programme in Leeds. Teachers distributed information during scheduled class sessions, and students were invited to participate on a voluntary basis. School-level agreement was obtained, and student assent was recorded prior to participation in accordance with institutional ethics approval. Participation was not mandatory, students could withdraw at any time without consequence, and no financial or other incentives were provided. Workshops for both studies were delivered during school enrichment activities or timetabled lessons.

In the first study, seven workshops were undertaken with 79 secondary school students across four schools, where students designed their own neuroimaging studies and identified priorities for research. In the second study, ten priorities derived from previous workshops and 376 students from four different schools were asked to rank these priorities from most important to least important. The two-stage design created a feedback process in which themes generated in participatory workshops were subsequently evaluated within a larger student sample. The data collected were analysed to identify common themes and priorities among the students. Deprivation profiles of participating schools were mapped using the Indices of Multiple Deprivation (IMD), as shown in Figure 1.

**Figure 1:**
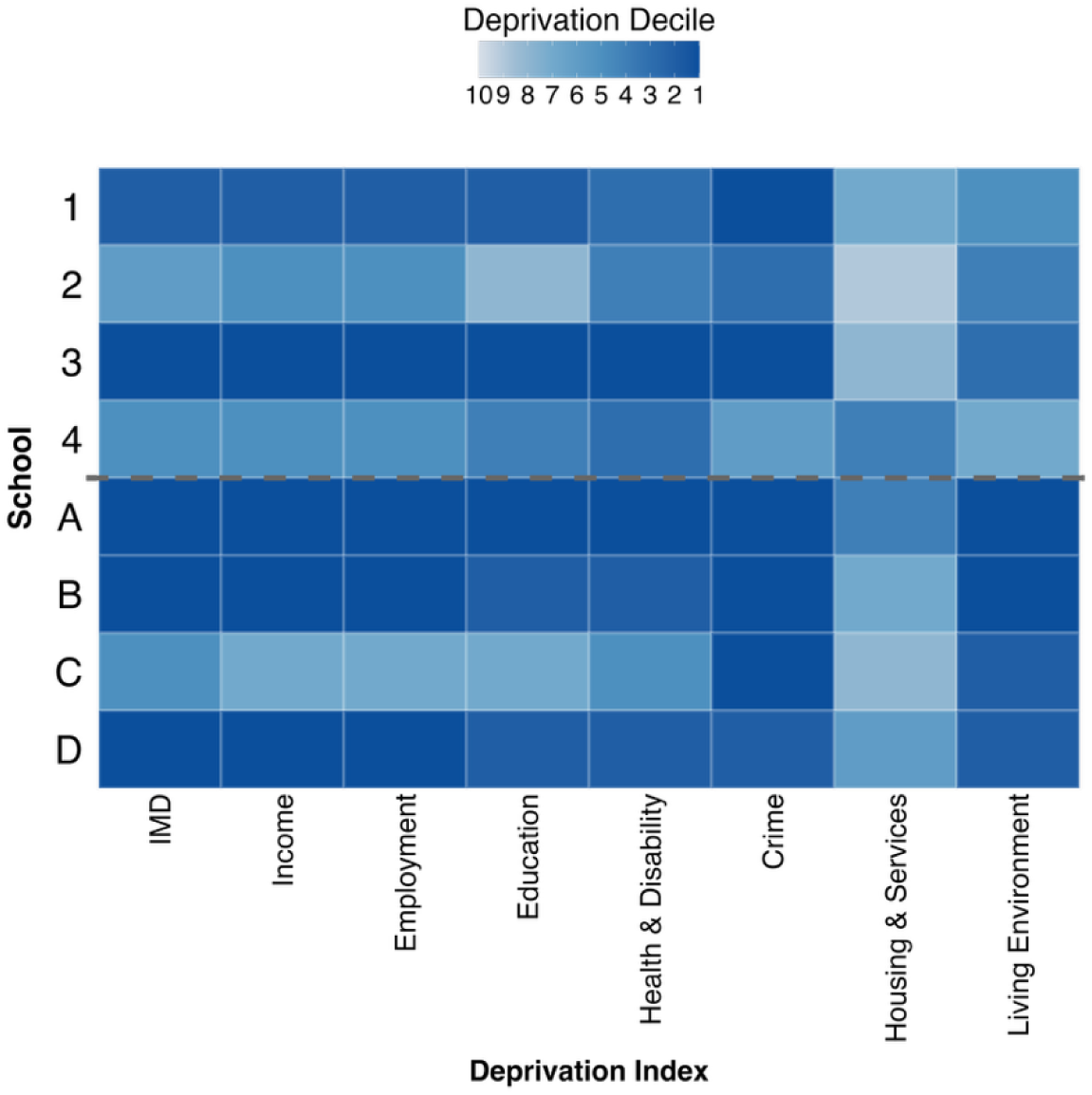
Indices of Multiple Deprivation for participating schools in Study 1 (1, 2, 3, 4), generating priorities and Study 2 (A, B, C, D), ranking priorities. Index deciles are presented for overall Index of Multiple Deprivation (IMD), Income, Employment, Education, Health and Disability, Crime, Barriers to Housing and Services and Living Environment. Darker colours indicate higher deprivation decile (1 = most deprived, 10 = least deprived).

To mitigate the potential impact of power imbalances, facilitators explicitly positioned adolescents as experts of their own experiences, with researchers acting as collaborators rather than evaluators. Students were explicitly told their input would inform our research programme. After workshops, schools received feedback reports summarising findings, closing the participation loop.

### Operationalisation of Community-Based Participatory Research Principles

Several Community-Based Participatory Research (CBPR) principles were operationalised within this work (Israel et al., 1998; Wallerstein & Duran, 2010). First, adolescents were engaged as a relevant community whose perspectives were considered important for shaping research priorities in developmental neuroscience. Participatory workshops enabled students to generate and refine themes related to adolescent brain health and development, allowing their experiences and concerns to inform the direction of potential research questions. Second, workshop sessions incorporated an element of co-learning, where researchers introduced basic concepts of neuroscience and specifically, EEG, while students contributed insights into issues, they considered relevant to adolescent development. Third, the study design included an iterative process, whereby themes generated during participatory workshops (Study 1) were subsequently evaluated within a larger sample (Study 2).

## Study 1: Priority Setting

### Methodology

Workshops were conducted with students from School Years 7, 8, 9 and 12 (corresponding to ages 11-18 years) from four schools across the region.

Each workshop session lasted one hour and was structured into three components. The workshop began with a brief introduction to the fundamentals of neuroscience and neuroimaging, aimed at equipping students with the foundational knowledge needed for meaningful participation in the subsequent activities. This segment was integrated to promote participants’ self-efficacy and self-advocacy (Smith et al., 2021) in the session. Within this introductory segment, students were introduced to the plans for a new local population neuroimaging programme, where the goal is to use EEG to explore adolescent brain development. Following the introduction, students were divided into groups to design a neuroimaging study. Students were informed that their ideas may be used to define the priorities of the research programme.

As the introduction focused on EEG as the neuroimaging technique to be employed in the forthcoming programme, students were encouraged to design the study integrating the fundamentals of EEG they had learned in the session with their personal interests. The workshop concluded with a “Dragon’s Den” style format, based on the popular UK TV show (analogous to Shark Tank in the US), where each group pitched their study designs to the class through a short presentation. Students explained their research ideas, justified their methodologies, and responded to questions from peers and facilitators, providing a platform for critical feedback and discussion.

Following the presentation, students worked in small groups, determined by classroom logistics, to design a poster presenting their research ideas. They were asked to consider the question: “If you were a scientist studying the teenage brain using EEG, what would you study?” Supporting prompts were provided where needed, including: “What things do you think play a role in how your brain changes as you grow up?” and “How could we potentially use EEG to study this?” No specific design templates were given beyond the requirement to include a title, identify the problem to be addressed, propose an EEG-based research solution, and explain the societal benefits of the work.

Students used materials available within the school, primarily pens and paper, although one school, with access to a computer lab, produced digital posters that were subsequently printed. Posters in teaching and learning contexts provide insight into how students prioritise information (Lane, 2001), revealing what they consider most important through choices such as colour, text formatting, and sectioning. Research has shown that the process of creating a poster can foster independence (Mackey, 2004) and offers an effective medium for students who may struggle to articulate complex scientific concepts verbally or in writing. In these cases, posters enable students to represent their ideas visually, supporting the development of critical thinking and enhancing both comprehension and communication of challenging material (Hasio, 2015). As such, the poster task provided a structured but flexible means of eliciting student priorities.

## Results

### Data Preparation and Analysis

Posters created by students were scanned and uploaded to NVivo 14 (Lumivero, 2023). Reflexive Thematic Analysis, following Braun & Clarke (2019), was applied to code generation of visual files. Disagreements were resolved through discussion rather than consensus coding to preserve interpretative depth.

Reflexivity was considered throughout, with coders documenting their own positionality (e.g. disciplinary background, prior experience with adolescent co-research). Coding was conducted by authors KH and LK. KH is white female PhD student. LK is a post-doctoral researcher identifying as a British female of Moroccan and English background. We note this because these positionalities are relevant to the analytic process, as both authors bring prior experience in developmental neuroscience and adolescent co-research, alongside distinct cultural and disciplinary perspectives that could shape interpretation. Reflexive engagement with these perspectives was maintained throughout coding to support transparency in how themes were constructed from participants’ contributions. The research team involved in data acquisition comprised developmental neuroscience researchers with experience conducting school-based neuroimaging research and experience of delivering teaching sessions to participants aged 11-18 years. The wider project team comprised individuals with experience of implementation of adolescent developmental research and co-production in school environments.

Figure 2 shows the findings from the reflexive thematic analysis, providing a conceptual representation of primary themes, sub-themes, and their interrelationships. These themes: Health and Well-Being, Anti-Social Behaviour, Everyday Routines, and Social Media Use, reflect our participants interests in understanding the neural mechanisms underlying their daily experiences, concerns, and behaviours. Within each theme, students identified specific sub-themes, highlighting areas where they believed neuroscience research could provide valuable insights. We explore these subthemes next.

**Figure 2:**
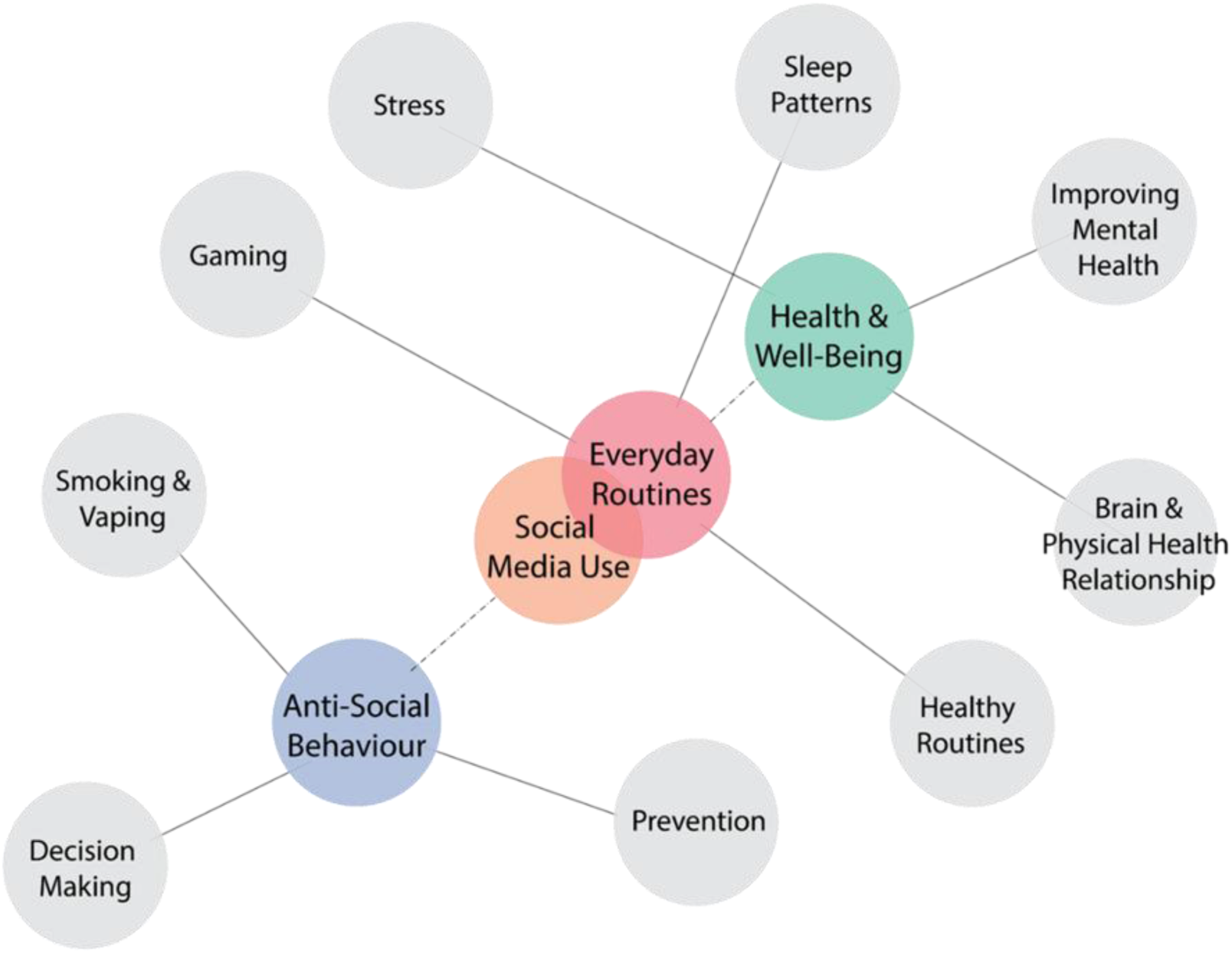
Thematic map derived from reflexive thematic analysis of the workshop posters. The map preserves participant-generated labels where possible and shows four overarching themes (in coloured circles and linked with dotted lines) alongside connected subthemes (in solid lines). Overlap indicates conceptual proximity within the analytic map.

### Theme: Health and Well-Being

Participants conceptualised well-being as a broad construct incorporating mental health, stress, and physical health. Their posters reflected a keen interest in the relationship between brain activity and external stressors, emotions, and lifestyle factors.

Stress emerged as a prominent sub-theme, with particular emphasis on exam-related stress. Students were concerned about the impact of academic pressure on brain function and proposed research comparing brain activity across different exam conditions. They suggested exploring whether stress responses vary depending on age and academic stage. Additionally, they speculated that neuroimaging techniques, such as EEG, could be used to identify unconscious stress triggers in students who may struggle to articulate their experiences.

Beyond academic stress, students considered the influence of social perception on brain activity. They hypothesised that external factors, such as clothing and social environments, could shape students’ self-perception and stress responses. Some proposed comparing brain activity in different social contexts to investigate whether these effects vary by gender or age group. They saw potential for such research to inform the development of supportive school environments that mitigate stress.

Mental health was a major concern, with students expressing a desire to understand the neural basis of conditions such as anxiety, depression, and phobias. Many saw neuroimaging as a tool to uncover how emotional states are represented in the brain. Some extended this interest to broader societal issues, proposing research into the neural mechanisms of racial bias and prejudice. They speculated that identifying patterns of brain activity associated with bias could help inform strategies for reducing discrimination.

Physical health was another key area of interest, particularly in relation to exercise and its impact on brain activity. Students proposed comparing brain responses across different exercise types, fitness levels, and time points surrounding physical activity. Some explored the idea that physical activity could interact with factors such as energy levels and stress, influencing cognitive function. They were particularly interested in how these relationships might vary by age and whether neuroscience could offer evidence-based insights into the benefits of exercise for brain health.

### Theme: Investigating Anti-Social Behaviour

Concerns about risk-taking behaviours and their neural underpinnings were central to this theme. Adolescents focused on decision-making processes, substance use, and the role of social influence in shaping behaviour.

Decision-making was identified as a critical developmental process. Students proposed creating controlled environments in which adolescents are presented with moral dilemmas or risky choices while their brain activity is monitored. They proposed this could provide insight into impulsivity and help develop interventions to support better decision-making. Some extended this interest to criminal behaviour, suggesting that research could explore whether certain brain activity patterns are associated with criminal tendencies.

Social influence was another key consideration. Adolescents were particularly interested in how peer pressure affects decision-making and morality. They hypothesised that social environments shape neural responses and suggested using neuroimaging to examine how individuals respond to social pressure when making choices.

Substance use, particularly smoking and vaping, emerged as another major concern. Students were curious about the impact of nicotine and other substances on adolescent brain development. They speculated that EEG could be used to investigate the neural mechanisms of addiction and to identify potential intervention strategies. Some extended this interest to broader issues of substance abuse, including alcohol and other drugs, proposing that neuroscience might offer insights into prevention and treatment.

### Theme: Everyday Routines

Participants also recognised the importance of structured routines in shaping cognitive and emotional well-being. Their posters reflected an interest in how everyday habits, such as sleep and gaming, influence brain function.

Sleep was a particularly prominent sub-theme. Students wanted to investigate the factors that affect sleep quality, duration, and latency, considering environmental variables such as temperature, noise, and bedding. Many proposed comparing the sleep patterns of different individuals to identify best practices for improving sleep. Some also speculated about the potential of neuroimaging to visualise dreams, questioning whether brain activity patterns could be used to illustrate dream content without verbal input.

Gaming was another area of interest, with students reflecting on their own experiences with video games and their potential effects on brain development. They hypothesised that different genres and design features, such as colours, haptics, and music, might have distinct effects on neural activity. Some were particularly interested in whether gaming influences mental health or contributes to antisocial behaviours, such as gaming-induced aggression.

### Theme: Social Media Use

Social media was recognised as both a distinct theme and a component of adolescents’ everyday routines. Students considered its impact on self-perception, mental health, and cognitive function.

Online interactions and self-perception were key concerns. Students discussed how social media influences their self-image, particularly in relation to gender differences. They speculated that exposure to social media might shape brain activity linked to self-perception and emotional responses. Some suggested that these effects might be moderated by factors such as screen time and age, warranting further investigation.

Mental health was also a central focus surrounding social media. Adolescents were concerned about the potential for social media to contribute to stress, anxiety, and mood fluctuations. They were particularly interested in whether neuroimaging could reveal patterns of brain activity associated with digital well-being.

### Thematic Intersections

Beyond overlaps within themes, several overlaps were evident, highlighting the interconnected nature of adolescents’ concerns. The intersection between gaming and mental health was a recurring topic, with students questioning whether video games contribute to emotional regulation difficulties or antisocial behaviours such as aggression. Similarly, stress and sleep were frequently discussed together, as students recognised that academic stress could disrupt sleep patterns and affect cognitive function.

Another notable intersection was between social media use and self-perception, where students hypothesised that online interactions influence stress, decision-making, and broader mental health outcomes. They suggested that these relationships might be particularly pronounced during adolescence, given the importance of peer validation.

## Study 1 Discussion

Mental health emerged as a consistently significant theme, highlighting its universal importance across different school environments and demographic groups. Adolescence is a critical period for mental health development, with stress playing a central role in shaping brain structures such as the hypothalamic-pituitary-adrenal axis, amygdala, and prefrontal cortex (Casey et al., 2018; Lupien et al., 2009). Persistent stress can interfere with these processes, leading to difficulties in emotional regulation and decision-making (Paus et al., 2008). Electroencephalography (EEG) has been shown to provide insights into the neural mechanisms underlying mental health conditions such as anxiety and depression (Boby & Veerasingam, 2025; Elnaggar et al., 2025; Gkintoni et al., 2025). Understanding these mechanisms might inform targeted interventions to improve adolescent well-being.

Stress and physical health were also ranked as important themes, whilst revealing some variation across schools and gender groups. This suggests that while these concerns are widely recognised, their perceived relevance may be shaped by local contexts and demographic factors. Research indicates that chronic stress can have lasting effects on neurodevelopment, particularly in areas governing executive function and emotional processing (Suleiman & Dahl, 2017). Similarly, disruptions to sleep-another key concern raised by students-have been shown to negatively impact cognitive, emotional, and physical well-being (Born in Bradford, 2024). Adolescents are particularly vulnerable to sleep deprivation due to biological shifts in circadian rhythms, compounded by academic and social pressures (Beebe, 2011).

Anti-Social Behaviour emerged as a distinct theme. Students highlighted the behavioural implications of excessive digital media consumption, linking it to reduced face-to-face interactions and lower social empathy. The scientific literature supports these observations, associating heightened digital engagement with social withdrawal and decreased interpersonal sensitivity (Twenge et al., 2018). Similarly, social media use was highlighted as both a standalone theme and one intersecting with everyday routines. Students acknowledged its substantial influence on their daily schedules, decision-making processes, and peer interactions. There is a growing body of research showing that excessive social media use can disrupt essential routines, impair cognitive function, and contribute to emotional instability (Orben & Przybylski, 2019). Furthermore, social media has been linked to anti-social behaviours, including cyberbullying and social withdrawal. The dual categorisation of social media illustrates its role as both an essential daily activity and a source of socio-behavioural challenges.

The theme of “Everyday Routines” surfaced prominently in students’ reflections, with subthemes such as Gaming and Sleep Patterns frequently mentioned. Students described gaming as an integral activity influencing time management, social engagement, and concentration. However, research suggests that excessive gaming can also precipitate anti-social behaviour by reducing real-world social interactions (Przybylski & Weinstein, 2017), though potentially moderated by the type of game (Greitemeyer, 2022).

Participants frequently presented ideas spanning multiple themes and sub-themes, demonstrating an awareness of the complexities of development and the interconnected nature of cognitive, emotional, and social processes. Their proposals often linked mental health to sleep, stress to decision-making, and gaming to social behaviour, reflecting an intuitive understanding that brain development does not occur in isolation. This overlap suggests a strong curiosity about how different aspects of their experiences interact at a neural level, highlighting their interest in exploring multifaceted relationships between behaviour, environment, and brain function.

## Study 2: Priority Ranking

Study 2 was designed as a validation phase to assess the relative importance of themes generated during the participatory workshops in Study 1 across a larger sample. Themes generated in Study 1 were translated into ten survey items for Study 2, which participants were asked to rank in order of importance. The broad health and well-being theme was disaggregated into mental health, physical health, and stress to provide greater granularity.

## Methodology

### Participants

A total of 376 participants were included in Study 2. These participants were distinct from those involved in Study 1 to reduce potential bias in the ranking task and to strengthen the validity of the priority-setting process. We reasoned that including the same participants could have led individuals to favour themes they had personally contributed during the workshops, which would have conflated idea generation with evaluation.

Demographic data were partially missing, with valid information available for 325 participants for Gender, 332 for Ethnicity, and 297 for Age. Missingness primarily reflected variations in session formats, where demographic data were not consistently collected, alongside participants’ right to withhold this information. Based on available data, the mean age of participants was 16.85 years (SD = 1.97, n = 297). Reported gender was predominantly female (n = 197) and male (n = 122), with one participant identifying as non-binary/third gender and five opting not to disclose.

The largest ethnic group identified as English, Welsh, Scottish, Northern Irish or British (*n* = 155). Other reported ethnicities included Pakistani (*n* = 51), Indian (*n* = 23), African (*n* = 19), any other White background (*n* = 28), “any other Black, Black British, or Caribbean background” (*n* = 11), White and Asian (*n* = 9), “any other Asian background” (*n* = 8), Chinese (*n* = 3), White and Black African (*n* = 3), White and Black Caribbean (*n* = 2), Caribbean (*n* = 2), Arab (*n* = 2), Irish (*n* = 2), Bangladeshi (*n* = 1), and “any other ethnic group” (*n* = 1). A further seven participants described mixed or multiple backgrounds across variants, and five selected “prefer not to say.”

The participants came from four schools (A, B, C, D in Figure 1) in Bradford and Leeds, though demographic data for School A was unavailable, due to its private, independent status. Socioeconomic indicators varied across schools. School B had the highest proportion of students eligible for free school meals (over 45%), exceeding both School C and the national mainstream average (approximately 25%). School B also reported the highest rate of student absences (nearly 20%), while School C had a lower absence rate but remained above the national average. Academic attainment, measured by the percentage of students achieving Grade 5 or above in key subjects, was highest in the national mainstream average, followed by School C, with School B showing the lowest performance.

### Approach

Sessions were delivered within classroom environments for a duration of a timetabled lesson (∼1 hour). Each session followed a structured format. In the first segment of the session, students received a brief introduction to the basic concepts of neuroscience. This included an overview of brain structure and function, as well as the relevance of neuroimaging in understanding brain activity. Following the introduction, students were given a demonstration of electroencephalography (EEG) technology. This demonstration included an explanation of how EEG works, its applications in research, and a live demonstration of EEG data collection. After the demonstration, students were presented with a list of ten research priorities that had been extracted from themes identified in Study 1: Mental Health, Stress, Relationship between Brain and Physical Health, Decision Making, Anti-Social Behaviour, Social Media, Healthy Routines, Gaming, Sleep, Smoking and Vaping.

The decision to split the broad ‘Health and Wellbeing’ theme into subdomains (mental health, physical health, stress) was based on its high prevalence in Study 1 and our intention to capture the more granular nuances adolescents had highlighted through their posters. This subdivision enabled a more precise analysis of how health-related priorities were ranked. The remaining themes were held constant across participants because allowing modification at this stage would have reduced comparability and limited the ability to assess the relative salience of themes generated in Study 1.

### Analysis

Responses were collected through a survey administered on paper or online via the Qualtrics platform. Participants were asked to rank a set of research priorities extracted from Study 1. Their responses, including demographic information such as gender and school postcode, were recorded and digitised. Data were aggregated to calculate the mean ranking for each research priority. This aggregation was performed both for the combined dataset of all participants (Figure 3). Visualisation of the aggregated rankings was conducted using R (4.4.1). The rankings were plotted from highest *(1)* to lowest *(10)*, illustrating the research priorities across all participants. As exploratory analysis, we examined whether priorities varied according to gender, age and school.

**Figure 3:**
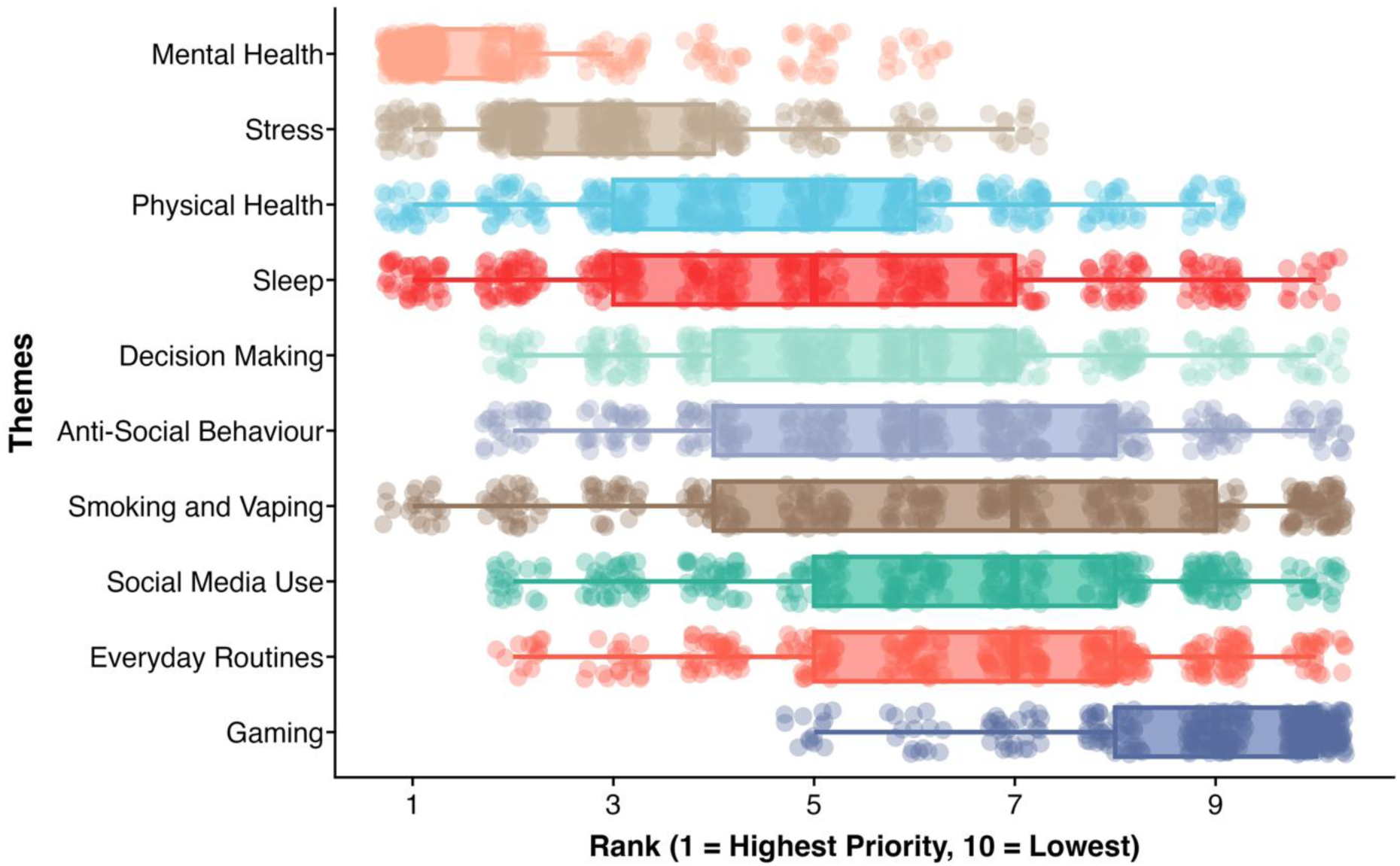
Distribution of average ranks across the ten priorities. Lower values indicate higher priority. Dots represent individual responses and boxplots summarise the central tendency and spread of ranks

**Figure 4:**
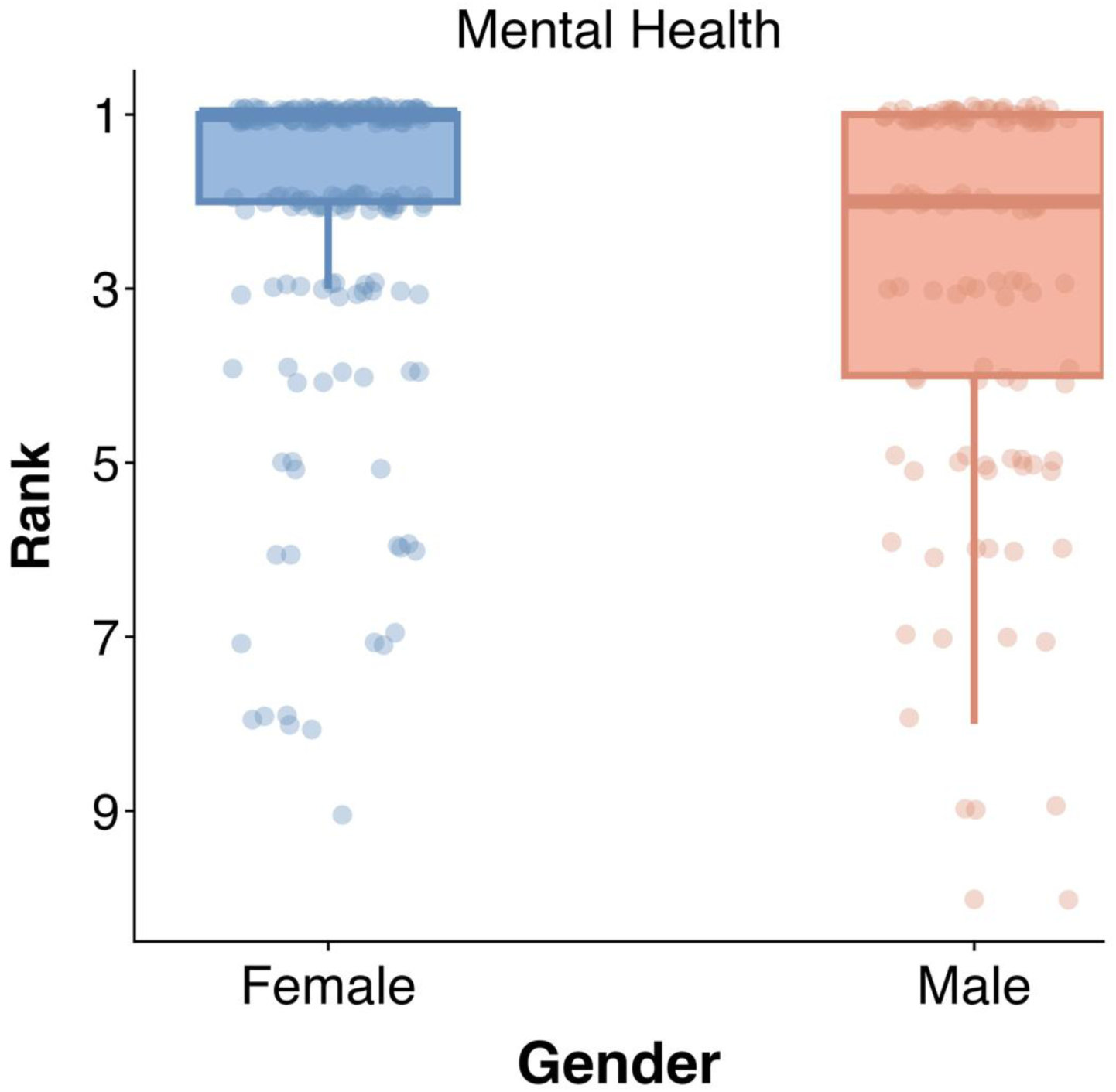
Ranked importance of mental health by gender. Lower values indicate higher priority. The figure shows the only gender difference that remained significant after FDR correction.

**Figure 5:**
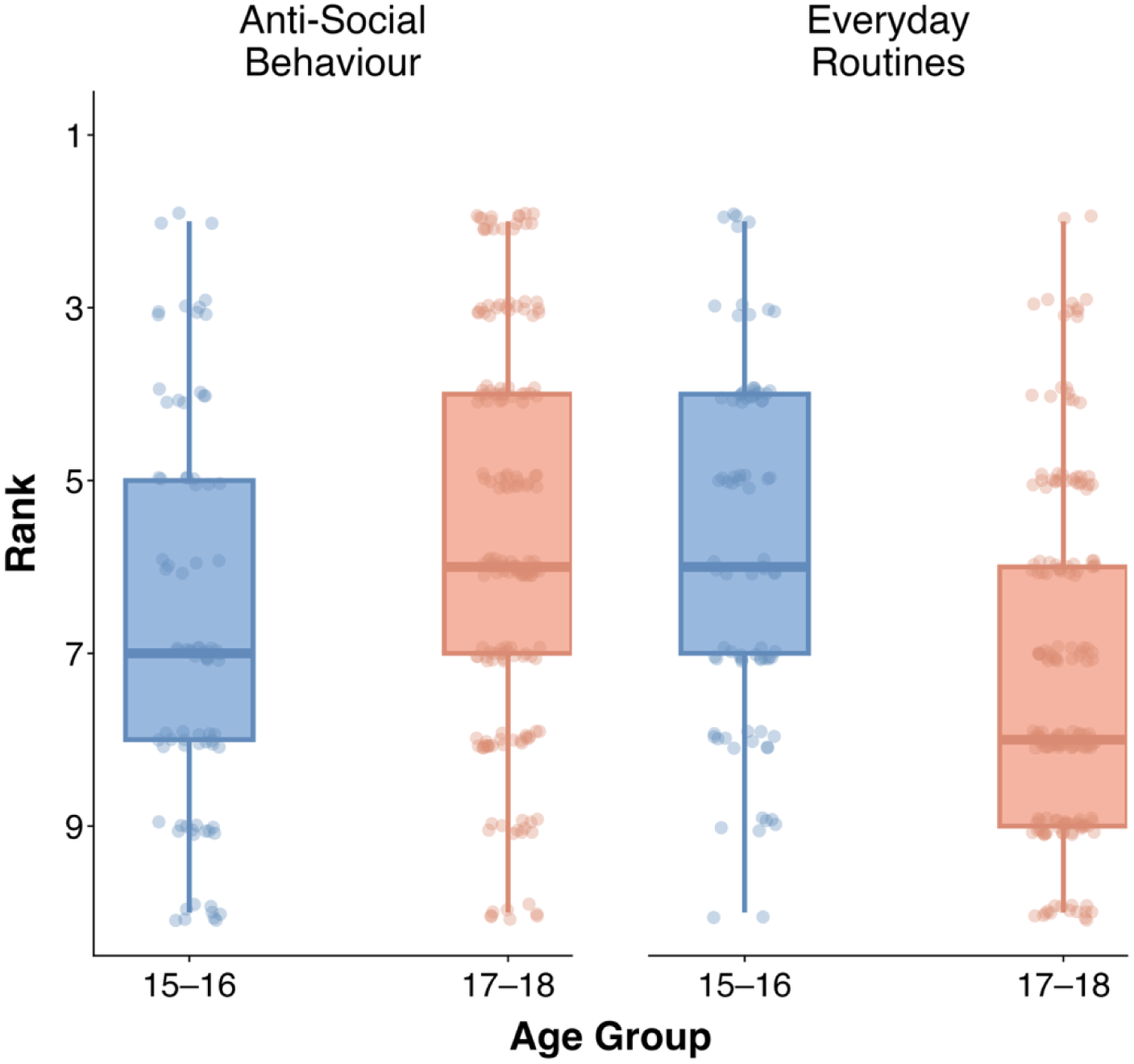
Age-related differences in priorities for the two themes that showed statistically significant differences. Older adolescents prioritised the study of anti-social behaviour while younger participants ranked the theme of “everyday routines” more highly. Lower values indicate higher priority.

**Figure 6:**
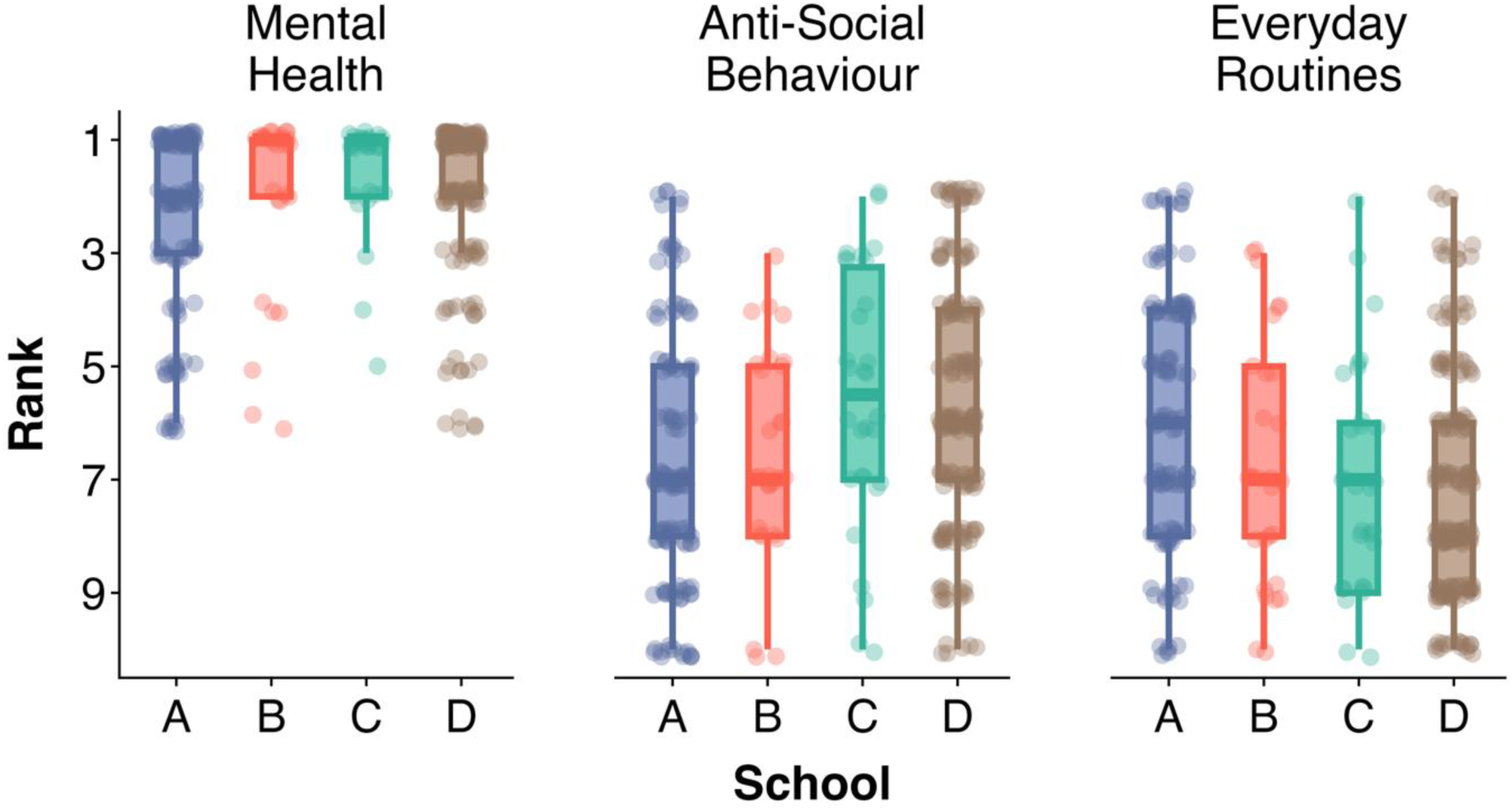
Differences in priorities across schools for themes Mental Health, Anti-Social Behaviour and Everyday Routines. Lower values indicate higher priority.

## Results

### Average Ranking

On average, participants ranked Mental Health as the highest priority (M = 2.39, SD = 2.08, *n* = 370), followed by Stress (M = 3.49, SD = 2.02) and Sleep (M = 4.90, SD = 2.72). Physical Health (M = 5.00, SD = 2.39), Decision Making (M = 5.59, SD = 2.31), and Anti-Social Behaviour (M = 5.93, SD = 2.41) were rated at an intermediate level of priority. Lower priority themes included Social Media Use (M = 6.25, SD = 2.47), Smoking and Vaping (M = 6.34, SD = 2.87), Everyday Routines (M = 6.51, SD = 2.34), and Gaming (M = 8.48, SD = 2.11).

### Rankings by Gender

We conducted exploratory analysis to examine whether prioritisation varied for gender. Mann–Whitney U tests with FDR correction showed that females placed significantly greater emphasis on Mental Health (U = 9601.5, p = .045, FDR-corrected), compared to males (M = 2.06, SD = 1.79, n = 194 vs. M = 2.80, SD = 2.31, n = 120). The associated effect size was small (rank-biserial r = .18; Cliff’s δ = −.18), indicating modest gender differences. No other themes survived FDR correction. Although raw p-values suggested possible differences for Sleep (p = .021), Physical Health, (p = .049), Smoking and Vaping (p = .026), Social Media Use (p = .019), and Gaming (p = .033), these did not remain significant after FDR adjustment (all p’s > .05). Thus, after controlling for multiple comparisons, gender-related differences were restricted only to the prioritisation of Mental Health, with all other themes showing negligible to small differences.

### Rankings by Age

To further explore potential differences in student rankings, we undertook exploratory analysis of age, splitting our participants into mid-adolescents (15–16 years) and late adolescents (17–18 years). Mann–Whitney U tests with FDR correction showed that late adolescents (M = 5.69, SD = 2.43, n = 179) placed significantly greater emphasis on Anti-Social Behaviour (U = 5928, p < .001, FDR-corrected), compared to mid-adolescents (M = 6.73, SD = 2.28, n = 83). The associated effect size was small (r = .20), indicating modest age-related differences. In contrast, mid-adolescents placed significantly greater emphasis on Everyday Routines (U = 1136, p < .001, FDR-corrected), compared to late adolescents (M = 5.76, SD = 2.41, n = 85 vs. M = 7.15, SD = 2.15, n = 182), with a large effect size (rank-biserial r = .85; Cliff’s δ = −.85). No other themes survived FDR correction. Although raw p-values suggested a possible difference for Mental Health (p = .001), this did not remain significant after FDR adjustment in the filtered analysis (p = .077). Thus, after controlling for multiple comparisons, age-related differences were restricted to Anti-Social Behaviour and Everyday Routines.

### Priorities by School

A series of Kruskal–Wallis tests were conducted to examine differences in ranked research priorities across four schools (A–D). Significant effects were observed for Mental Health, χ²(3) = 14.66, p = .007, ε² = .03; Anti-Social Behaviour, χ²(3) = 18.90, p = .001, ε² = .05; and Everyday Routines, χ²(3) = 25.83, p < .001, ε² = .06.

No significant differences were found for Stress, Physical Health, Sleep, Decision Making, Smoking and Vaping, Social Media Use, or Gaming (all *p*’s > .05). The observed effect sizes were small in magnitude, with no theme exceeding the threshold for a medium effect (ε² ≥ .06).

Follow-up pairwise Dunn tests with Benjamini–Hochberg adjustment showed the following significant contrasts: Mental Health was ranked highest priority in School C (M = 1.54, SD = 1.00, n = 28) compared to School A (M = 2.39, SD = 1.63, n = 114), School B (M = 2.04, SD = 1.63, n = 27), and School D (M = 1.80, SD = 1.31, n = 177). Anti-Social Behaviour was given greater importance in School C (M = 5.53, SD = 2.29, n = 30) and School D (M = 5.74, SD = 2.18, n = 178) compared with School A (M = 6.75, SD = 2.28, n = 120). Everyday Routines was prioritised lowest in School D (M = 7.16, SD = 2.01, n = 180) relative to School A (M = 5.88, SD = 2.22, n = 121) and School B (M = 6.69, SD = 2.14, n = 29). No other pairwise comparisons were significant after FDR correction.

## General Discussion

We employed a two-stage participatory approach to identify adolescents’ priorities for developmental neuroimaging research. Four broad themes emerged: mental health and wellbeing, antisocial behaviour, everyday routines, and social media. From these themes, mental health consistently ranked as the highest priority across schools, genders and age groups. Differences between genders were modest, with female participants emphasising mental health more strongly. Differences were also observed across age groups and schools, particularly for antisocial behaviour and everyday routines. These contextual variations suggest that priorities may be shaped not only by individual characteristics but also by the environments in which adolescents live and learn.

We note that “Health and Well-Being” featured prominently in the posters developed in Study 1. Given the prevalence of this broad theme, we sought to explore this in more detail and as such, separated this category into three: “Mental Health”, “Physical Health” and “Stress” for Study 2. We considered whether the high prioritisation of mental health reflected its broad and multi-faceted nature, potentially encompassing related domains such as stress, sleep, and social media use. While this is a plausible interpretation, adolescents in the present study consistently generated mental health alongside these domains as distinct areas, often describing them independently within their proposed research ideas. Mental health was therefore retained as a subtheme within the broader “health and well-being” category to reflect the structure of participants’ contributions, rather than imposing researcher-defined hierarchies. This approach is consistent with participatory methodologies that prioritise participant-defined structures (Clark et al., 2022; Mikesell et al., 2013). It also reflects evidence that adolescent perspectives may not map directly onto established theoretical constructs, but instead capture experiences in ways that are meaningful within their own contexts (Fuligni, 2019; Toenders et al., 2024). Future work could more explicitly examine how young people conceptualise the relationships between these domains.

In our exploratory analysis we found some differences according to participant demographics. Female students consistently prioritised mental health, mirroring findings from previous community-based research (Born in Bradford, 2024), while other priorities remained broadly consistent across genders. Age-related analyses suggested that research priorities were largely stable across mid-to-late adolescence, with some specific shifts: Younger adolescents placed greater emphasis on everyday routines, whereas older adolescents prioritised anti-social behaviour more highly. One possible interpretation is that these patterns reflect developmental differences in daily structure, autonomy, and social context (Salmela-Aro, 2011). Younger adolescents are typically more embedded in structured routines coordinated through caregivers and schools, while older adolescents experience increased independence and greater exposure to complex social and behavioural dynamics, which may heighten the salience of anti-social behaviour as a concern. Adolescence is also characterised by continued maturation of prefrontal regions alongside heightened sensitivity to social contexts (Casey et al., 2018; Steinberg, 2008). As such, variation in the prioritisation of antisocial behaviour may not only reflect contextual factors, but also developmental differences in how these behaviours are perceived, regulated, and understood. However, the present study was not designed to examine how priorities vary across the full developmental trajectory. It is not clear whether these patterns would generalise across a broader developmental range, including earlier adolescence or the transition into adulthood, where further changes in autonomy, identity, and social roles occur.

The present work also highlights the importance of situating neuroscience, and health research more broadly, within the contexts in which data are generated. In many areas of health and neuroimaging research, race and ethnicity have historically been included in statistical models as broad demographic variables, sometimes standing in for unmeasured socioeconomic or environmental factors. However, there is increasing recognition that this practice can obscure the underlying structural drivers of observed differences and risks reinforcing misleading interpretations (Cardenas-Iniguez et al., 2024). This has led to calls for greater emphasis on context-sensitive measures that more directly capture environmental and structural variability (Cardenas-Iniguez et al., 2024). One such approach is the use of socioeconomic and deprivation indices to characterise the environments in which participants are embedded. Adopting a “place-based” approach, situating data collection and analysis within the specific cultural, socioeconomic, and environmental contexts of a defined geographic setting, acknowledging that local conditions significantly influence outcomes (Cummins et al., 2007) could help us move beyond broad demographic categories and attend more directly to the social and environmental conditions in which development unfolds.

In the present study, school-level deprivation profiles were used to contextualise differences observed across sites. While these differences were statistically small, they nevertheless suggest that adolescent priorities are not context-free. We found, for example, that students in School A, a private fee-paying institution, placed significantly lower priority on mental health compared to other schools. One possible explanation is differential access to institutional resources, as more resource-rich settings may provide greater support for mental health needs, potentially reducing the perceived urgency of this issue (DeAngelis & Dills, 2021). However, such interpretations are speculative, and causality cannot be inferred from the present data. Future work could test these hypotheses using designs that directly sample across socioeconomic contexts.

Participatory priority-setting is widely used in public health research but remains relatively uncommon in developmental cognitive neuroscience and adolescent mental health research (Norton, 2021). Many of the themes identified by adolescents in the present study align with areas already receiving attention in adolescent research, including mental health, sleep, stress, and digital media use. However, while these domains are already studied within developmental cognitive neuroscience, their operationalisation is typically researcher-defined. This raises the possibility that experimental paradigms capture constructs in ways that do not fully reflect how adolescents experience or interpret them. Participatory priority-setting therefore represents an important first step, not in redefining what is studied, but in refining how these constructs are operationalised and prioritised within experimental design. For example, emphasis on context-specific stressors, such as academic pressure and social evaluation, suggests a potential need for more ecologically valid paradigms that incorporate meaningful real-world conditions, rather than abstract task designs. Similarly, the prioritisation of sleep, social media use, and everyday routines highlights the importance of integrating behavioural and environmental measures alongside neural data, ensuring that these factors are treated as central components of study design rather than secondary covariates. In this way, participatory approaches could inform not only which constructs are studied, but how they are operationalised within broader experimental designs.

The co-design process in the present study focused on identifying research priorities rather than collaboratively developing experimental paradigms. While students were introduced to basic neuroscience concepts, designing experimentally valid neuroimaging tasks requires methodological expertise that cannot be developed within a short workshop setting. Adolescents were engaged intensively in the generation and prioritisation of research questions, but they did not take part in data analysis, nor in governance or authorship decisions. Given logistical constraints, we focussed on a staged approach consistent with CBPR, in which early phases prioritise agenda-setting before progressing to the co-development of research tools (Israel et al., 1998; Wallerstein & Duran, 2010). The present study lays the groundwork for future work in which selected priorities can be taken forward into the collaborative development of neuroimaging paradigms, enabling a more reciprocal approach to defining and operationalising constructs within developmental cognitive neuroscience (La Scala et al., 2023).

Finally, the positionality of the research team is relevant when it comes to interpreting these findings. As developmental neuroscience researchers with experience conducting school-based neuroimaging studies, we are regularly exposed to the questions and curiosities young people express when engaging with neuroscience research. This familiarity likely supported our interpretation of the workshop discussions and situating of themes within developmental neuroscience. At the same time, our experience within the field may have influenced how participants’ ideas were organised and translated into research-relevant constructs. While this interpretive influence should be acknowledged, it is also an inherent part of translating community perspectives into research questions that can be addressed within developmental neuroscience.

## Conclusions

This study demonstrates the value of involving adolescents in setting priorities for developmental neuroscience. We argue that participatory priority-setting should become a standard early step in developmental neuroimaging programmes, especially those seeking translational or policy relevance. Doing so can help ensure that future research agendas reflect not only scientific opportunity, but also adolescent concerns, context, and lived experience.

## Contributions

**Conceptualisation:** Katherine Hiley, Layla Kouara, Faisal Mushtaq. **Data curation:** Katherine Hiley, Layla Kouara, David Ryan, Katy Shire, Zarina Mirza. **Formal analysis:** Katherine Hiley, Layla Kouara. **Funding acquisition:** Faisal Mushtaq, Layla Kouara. **Investigation:** Katherine Hiley, Layla Kouara. **Methodology:** Katherine Hiley, Layla Kouara. **Project administration:** Katherine Hiley. **Resources:** Katherine Hiley and Layla Kouara. **Supervision:** Faisal Mushtaq. **Validation:** Faisal Mushtaq. **Visualisation:** Katherine Hiley, Layla Kouara and Faisal Mushtaq. **Writing - original draft:** Katherine Hiley, Layla Kouara, and Faisal Mushtaq. **Writing - review & editing:** Katherine Hiley, Layla Kouara, David Ryan, Katy Shire, Zarina Mirza, Faisal Mushtaq.

## Acknowledgements

This project was made possible thanks to a Participatory Research Fund award from Research England. KH’s PhD was funded in part by the National Institute for Health and Care Research Yorkshire and Humber ARC (NIHR200166) and the University of Leeds. LK was supported by an Emma and Leslie Reid PhD Scholarship. FM is supported by the UK Research and Innovation Biotechnology and Biological Sciences Research Council (BB/X008428/1) and a Special Projects award from the Huo Family Foundation. FM and MMW are both supported by the National Institute for Health and Care Research (NIHR) Leeds Biomedical Research Centre (NIHR203331). The views expressed are those of the authors and not necessarily those of the NIHR or the Department of Health and Social Care.

## Data Availability

All anonymised posters and raw ranking data generated during this study are publicly available on the Open Science Framework (OSF) repository. The data can be accessed at https://osf.io/f2eg3/?view_only=f7102e57a77f4ba9bd60a9659b163ddc.

## References

1. Abraczinskas, M., & Zarrett, N. (2020). Youth Participatory Action Research for Health Equity: Increasing Youth Empowerment and Decreasing Physical Activity Access Inequities in Under-resourced Programs and Schools. American Journal of Community Psychology, 66. 10.1002/ajcp.12433

2. Beebe, D. W. (2011). Cognitive, Behavioral, and Functional Consequences of Inadequate Sleep in Children and Adolescents. Pediatric Clinics of North America, 58(3), 649–665. 10.1016/j.pcl.2011.03.002

3. Boby, K., & Veerasingam, S. (2025). Depression diagnosis: EEG-based cognitive biomarkers and machine learning. Behavioural Brain Research, 478, 115325. 10.1016/j.bbr.2024.115325

4. Born in Bradford. (2024). Mental Health and Wellbeing in Schools. Born in Braford. https://borninbradford.nhs.uk/wp-content/uploads/2024/11/HG3165-BIHR-AoW-Mental-Health-and-Wellbeing-in-Schools4.pdf

5. Brady, L.-M., Miller, J., McFarlane-Rose, E., Noor, J., Noor, R., & Dahlmann-Noor, A. (2023). “We know that our voices are valued, and that people are actually going to listen”: Co-producing an evaluation of a young people’s research advisory group. Research Involvement and Engagement, 9(1), 11. 10.1186/s40900-023-00419-4

6. Cardenas-Iniguez, C., Schachner, J. N., Ip, K. I., Schertz, K. E., Gonzalez, M. R., Abad, S., & Herting, M. M. (2024). Building towards an adolescent neural urbanome: Expanding environmental measures using linked external data (LED) in the ABCD study. Developmental Cognitive Neuroscience, 65, 101338. 10.1016/j.dcn.2023.101338

7. Casey, B. J., Cannonier, T., Conley, M. I., Cohen, A. O., Barch, D. M., Heitzeg, M. M., Soules, M. E., Teslovich, T., Dellarco, D. V., Garavan, H., Orr, C. A., Wager, T. D., Banich, M. T., Speer, N. K., Sutherland, M. T., Riedel, M. C., Dick, A. S., Bjork, J. M., Thomas, K. M., … Dale, A. M. (2018). The Adolescent Brain Cognitive Development (ABCD) study: Imaging acquisition across 21 sites. *Developmental Cognitive Neuroscience, The Adolescent Brain Cognitive Development (ABCD) Consortium: Rationale*, Aims, and Assessment Strategy, 32, 43–54. 10.1016/j.dcn.2018.03.001

8. City of Bradford Metropolitan District Council. (2023). Bradford District Profile 2023.

9. Clark, A. T., Ahmed, I., Metzger, S., Walker, E., & Wylie, R. (2022). Moving From Co-Design to Co-Research: Engaging Youth Participation in Guided Qualitative Inquiry. International Journal of Qualitative Methods, 21, 16094069221084793. 10.1177/16094069221084793

10. Cummins, S., Curtis, S., Diez-Roux, A. V., & Macintyre, S. (2007). Understanding and representing ‘place’ in health research: A relational approach. Social Science & Medicine (1982), 65(9), 1825–1838. 10.1016/j.socscimed.2007.05.036

11. DeAngelis, C. A., & Dills, A. K. (2021). The effects of school choice on mental health. School Effectiveness and School Improvement, 32(2), 326–344. 10.1080/09243453.2020.1846569

12. Elnaggar, K., El-Gayar, M., & Elmogy, M. (2025). Depression Detection and Diagnosis Based on Electroencephalogram (EEG) Analysis: A Comprehensive Review. Diagnostics, 15(2), 210. 10.3390/diagnostics15020210

13. Farah, M. J. (2017). The Neuroscience of Socioeconomic Status: Correlates, Causes, and Consequences. Neuron, 96(1), 56–71. 10.1016/j.neuron.2017.08.034

14. Fuligni, A. J. (2019). The Need to Contribute During Adolescence. Perspectives on Psychological Science: A Journal of the Association for Psychological Science, 14(3), 331–343. 10.1177/1745691618805437

15. Gkintoni, E., Vantarakis, A., & Gourzis, P. (2025). Neuroimaging Insights into the Public Health Burden of Neuropsychiatric Disorders: A Systematic Review of Electroencephalography-Based Cognitive Biomarkers. Medicina, 61(6), 1003. 10.3390/medicina61061003

16. Greitemeyer, T. (2022). The dark and bright side of video game consumption: Effects of violent and prosocial video games. Current Opinion in Psychology, 46, 101326. 10.1016/j.copsyc.2022.101326

17. Hasio, C. (2015). Visual Inspirations: The Pedagogical and Cultural Significance of Creative Posters in the Art Classroom. Journal of College Teaching & Learning, 12(1), 39–44.

18. Israel, B. A., Schulz, A. J., Parker, E. A., & Becker, A. B. (1998). REVIEW OF COMMUNITY-BASED RESEARCH: Assessing Partnership Approaches to Improve Public Health. Annual Review of Public Health, 19(1), 173–202. 10.1146/annurev.publhealth.19.1.173

19. La Scala, S., Mullins, J. L., Firat, R. B., Emotional Learning Research Community Advisory Board, & Michalska, K. J. (2023). Equity, diversity, and inclusion in developmental neuroscience: Practical lessons from community-based participatory research. Frontiers in Integrative Neuroscience, 16. 10.3389/fnint.2022.1007249

20. Lane, T. K. (n.d.). Promoting Posters.

21. Lupien, S. J., McEwen, B. S., Gunnar, M. R., & Heim, C. (2009). Effects of stress throughout the lifespan on the brain, behaviour and cognition. Nature Reviews. Neuroscience, 10(6), 434–445. 10.1038/nrn2639

22. Mackey, A. (2004). TASK-BASED LANGUAGE LEARNING AND TEACHING. Studies in Second Language Acquisition, 26(3), 480–482. 10.1017/S0272263104293056

23. Marschall, M. (2004). Citizen Participation and the Neighborhood Context: A New Look at the Coproduction of Local Public Goods. Political Research Quarterly - POLIT RES QUART, 57. 10.2307/3219867

24. Mikesell, L., Bromley, E., & Khodyakov, D. (2013). Ethical Community-Engaged Research: A Literature Review. American Journal of Public Health, 103(12), e7–e14. 10.2105/AJPH.2013.301605

25. National Institute for Health and Care Research Global Health Research Unit on Global Surgery. (2023). Reducing the environmental impact of surgery on a global scale: Systematic review and co-prioritization with healthcare workers in 132 countries. The British Journal of Surgery, 110(7), 804–817. 10.1093/bjs/znad092

26. NIHR. (2024). NIHR Guidance on Co-Producing a Research Project. National Institute for Health Research (NIHR). https://www.learningforinvolvement.org.uk/content/resource/nihr-guidance-on-co-producing-a-research-project/

27. Norton, M. J. (2021). Co-Production within Child and Adolescent Mental Health: A Systematic Review. International Journal of Environmental Research and Public Health, 18(22), 11897. 10.3390/ijerph182211897

28. Orben, A., & Przybylski, A. K. (2019). The association between adolescent well-being and digital technology use. Nature Human Behaviour, 3(2), 173–182. 10.1038/s41562-018-0506-1

29. Ostrom, E. (1996). Crossing the great divide: Coproduction, synergy, and development. World Development, 24(6), 1073–1087. 10.1016/0305-750X(96)00023-X

30. Paus, T., Keshavan, M., & Giedd, J. N. (2008). Why do many psychiatric disorders emerge during adolescence? Nature Reviews. Neuroscience, 9(12), 947–957. 10.1038/nrn2513

31. Pavarini, G., Lorimer, J., Manzini, A., Goundrey-Smith, E., & Singh, I. (2019). Co-producing research with youth: The NeurOx young people’s advisory group model. Health Expectations, 22(4), 743–751. 10.1111/hex.12911

32. Pierce, M., Hope, H., Ford, T., Hatch, S., Hotopf, M., John, A., Kontopantelis, E., Webb, R., Wessely, S., McManus, S., & Abel, K. M. (2020). Mental health before and during the COVID-19 pandemic: A longitudinal probability sample survey of the UK population. The Lancet. Psychiatry, 7(10), 883–892. 10.1016/S2215-0366(20)30308-4

33. Przybylski, A. K., & Weinstein, N. (2017). A Large-Scale Test of the Goldilocks Hypothesis. Psychological Science, 28(2), 204–215. 10.1177/0956797616678438

34. Reiss, F. (2013). Socioeconomic inequalities and mental health problems in children and adolescents: A systematic review. Social Science & Medicine, 90, 24–31. 10.1016/j.socscimed.2013.04.026

35. Ricard, J. A., Parker, T. C., Dhamala, E., Kwasa, J., Allsop, A., & Holmes, A. J. (2023). Confronting racially exclusionary practices in the acquisition and analyses of neuroimaging data. Nature Neuroscience, 26(1), 4–11. 10.1038/s41593-022-01218-y

36. Ryan, D., Nutting, H., Parekh, C., Crookes, S., Southgate, L., Caines, K., Dear, P., John, A., Rehman, M. A., Davidson, D., Abid, U., Davidson, L., Shire, K. A., & McEachan, R. R. C. (2024). Ready, set, co(produce): A co-operative inquiry into co-producing research to explore adolescent health and wellbeing in the Born in Bradford Age of Wonder project. Research Involvement and Engagement, 10(1), 41. 10.1186/s40900-024-00578-y

37. Salmela-Aro, K. (2011). Stages of Adolescence. In B. B. Brown & M. J. Prinstein (Eds), Encyclopedia of Adolescence (pp. 360–368). Academic press.

38. Shire, K. A., Newsham, A., Rahman, A., Mason, D., Ryan, D., Lawlor, D. A., Opio-Te, G., Nutting, H., West, J., Pickavance, J., Dickerson, J., Pickett, K. E., Lennon, L., Gunning, L., Mon-Williams, M., Smith, S., Gilbody, S., Dogra, S., Walsh, T., … Wright, J. (2024). Born in Bradford’s Age of Wonder cohort: Protocol for adolescent data collection. Wellcome Open Research, 9, 32. 10.12688/wellcomeopenres.20785.1

39. Smith, H., Allf, B., Larson, L., Futch, S., Lundgren, L., Pacifici, L., & Cooper, C. (2021). Leveraging Citizen Science in a College Classroom to Build Interest and Efficacy for Science and the Environment. Citizen Science: Theory and Practice, 6(1), 29. 10.5334/cstp.434

40. Steinberg, L. (2008). A Social Neuroscience Perspective on Adolescent Risk-Taking. Developmental Review: DR, 28(1), 78–106. 10.1016/j.dr.2007.08.002

41. Suleiman, A. B., & Dahl, R. E. (2017). Leveraging Neuroscience to Inform Adolescent Health: The Need for an Innovative Transdisciplinary Developmental Science of Adolescence. Journal of Adolescent Health, 60(3), 240–248. 10.1016/j.jadohealth.2016.12.010

42. Tembo, D., Hickey, G., Montenegro, C., Chandler, D., Nelson, E., Porter, K., Dikomitis, L., Chambers, M., Chimbari, M., Mumba, N., Beresford, P., Ekiikina, P. O., Musesengwa, R., Staniszewska, S., Coldham, T., & Rennard, U. (2021). Effective engagement and involvement with community stakeholders in the co-production of global health research. The BMJ, 372, n178. 10.1136/bmj.n178

43. Thijssen, P., & Van Dooren, W. (2016). Who you are/where you live: Do neighbourhood characteristics explain co-production? International Review of Administrative Sciences, 82(1), 88–109. 10.1177/0020852315570554

44. Toenders, Y. J., Green, K. H., te Brinke, L. W., van der Cruijsen, R., van de Groep, S., & Crone, E. A. (2024). From developmental neuroscience to policy: A novel framework based on participatory research. Developmental Cognitive Neuroscience, 67, 101398. 10.1016/j.dcn.2024.101398

45. Twenge, J. M., Joiner, T. E., Rogers, M. L., & Martin, G. N. (2018). Increases in Depressive Symptoms, Suicide-Related Outcomes, and Suicide Rates Among U.S. Adolescents After 2010 and Links to Increased New Media Screen Time. Clinical Psychological Science, 6(1), 3–17. 10.1177/2167702617723376

46. Wallerstein, N., & Duran, B. (2010). Community-based participatory research contributions to intervention research: The intersection of science and practice to improve health equity. American Journal of Public Health, 100 Suppl 1(Suppl 1), S40–46. 10.2105/AJPH.2009.184036

47. Warraitch, A., Wacker, C., Biju, S., Lee, M., Bruce, D., Curran, P., Khraisha, Q., & Hadfield, K. (2024). Positive Impacts of Adolescent Involvement in Health Research: An Umbrella Review. Journal of Adolescent Health, 75(2), 218–230. 10.1016/j.jadohealth.2024.02.029

48. Webb, E. K., Etter, J. A., & Kwasa, J. A. (2022). Addressing racial and phenotypic bias in human neuroscience methods. Nature Neuroscience, 25(4), 410–414. 10.1038/s41593-022-01046-0

